# Use of *Ganoderma lucidum* grown on agricultural waste to remove antibiotics from water

**DOI:** 10.1101/2022.08.07.503092

**Authors:** Vanessa Salandez, Shiva Emami, Ameer Y. Taha, Valeria La Saponara

## Abstract

Antibiotic effluents from farming and medical applications into waterways pose serious risks for antibiotic drug resistance, promoting a need for effective strategies of removal from the environment. This experiment uses a novel mycoremediation approach to remove antibiotic contamination in synthetic wastewater. A white rot fungus, *Ganoderma lucidum*, was grown on biomass formed by agricultural waste from California (almond shells, fava bean stalks). Water containing or lacking *Ganoderma lucidum* was inoculated with twenty antibiotics from six different classes. The extent of antibiotic removal was measured at baseline and after 3 days with ultra-high pressure liquid chromatography coupled to tandem mass-spectrometry. In water containing *Ganoderma lucidum* mycelial biomass, we found a significant reduction compared to the baseline of the concentration in six (three quinolones and three sulfonamides) out of twenty tested antibiotics by Day 3, with normalized changes ranging from −24.4% to −82.4%. The mycelial biomass was particularly effective in reducing the presence of three quinolone antibiotics, a class of highly used antibiotics recalcitrant to processes in wastewater treatment plants. Our findings provide a novel approach to degrade certain antibiotics from water. This strategy could become a key component of removing antibiotic pollution using agricultural waste as part of the solution.

## 1. Introduction

Environmental pollution is an increasing global problem exacerbated by the rate of increasing population use of active pharmaceutical ingredients (API) such as antibiotics. ^[1]^ Antibiotics are used and misused in medical, veterinary, and farming (both aquaculture and agriculture) applications. They can enter the environment via wastewater effluents and agricultural runoffs, leading to the evolution and proliferation of antibiotic-resistant bacteria in wastewater, e.g. references. ^[2–8]^ Antibiotic contamination through excreted/secreted active components and metabolites can be harmful to human health and the ecosystem due to the spread of antibiotic resistant genes to humans and inhabitant species, e.g. references ^[4–5, 9]^, at very low concentrations. ^[10]^ Antibiotics can also impact organisms that are a vital part of the food chain, such as algae ^[4, 11]^ and cyanobacteria. ^[12]^

Several methods are currently used to remove antibiotics from wastewater, including activated carbon adsorption, membrane filtration, coagulation and flocculation, advanced oxidation processes, bioadsorption and activated sludge systems. ^[13]^ Unfortunately, the studies found in the literature so far show that these methods are not highly effective in removing antibiotics and vary in efficacy from plant to plant. Additionally, there are several studies reporting that the conditions in wastewater treatment plants are favorable to the proliferation of antibiotic-resistant bacteria. Wastewater and sewage sludge may be used as fertilizers, which means that they are constantly re-introduced to natural ecosystems, despite containing antibiotic-resistant bacteria. (reviews by Bouki et al. ^[14]^ and Fadário Frade et al. ^[11]^)

This paper focuses on a mycoremediation process, that is the degradation and removal of pollutants by filamentous fungi (also called bioremediation). Many filamentous fungi are able to degrade organic compounds present in vegetal and dead animals. The enzymatic system of fungi can also remove various toxic organic compounds, through sorption or degradation into less toxic/innocuous molecules, as observed in studies with hydrocarbons and polychlorinated biphenyls, PCBs. ^[15]^ Bioremediation has been successfully used since the 1980s (Bumpus et al. ^[16]^, researching the removal of DDT by *P. chrysosporium*).

On the other hand, mycoremediation studies aimed at degrading/reducing the concentration of antibiotics in contaminated water are fewer (review by Rodríguez-Rodríguez et al. ^[17]^). Studies under controlled conditions showed the ability of white rot fungi to transform antibiotics present in flasks of synthetic wastewater (aqueous solutions of antibiotics) or in bioreactors. The fungi are typically in the form of cultures grown in malt-based medium ^[13, 18–22]^ or in colonized agar. ^[17, 23]^ Several authors studied fungal pellets fed with malt extract agar in bioreactors treating wastewater sludge (e.g. references ^[24–26]^, and review by Espinosa-Ortiz et al. ^[27]^). We also make note of the study by Martens et al. ^[28]^, where the authors grew brown rot fungus (*Gloephyllum striatum*) in wheat straw or agricultural soil; the work of Chirnside and Kwart ^[29]^, on a white rot fungus (*Phanerochaete chrysosporium*) grown on corn stalks in a bioreactor for several weeks, to treat food processing wastewater; the experiments by Chakraborty and Abraham ^[30]^, where wood rotting fungus (*Ganoderma lucidum*) was isolated from hospital wastewater soil, and then maintained on potato dextrose agar plates.

In this paper, the mycelium (the root structure) of a white rot fungus, *Ganoderma lucidum*, was grown to investigate its remediation potential in synthetic wastewater while repurposing agricultural waste from California (USA). Agricultural waste biomass (e.g. hemp hurds, rice husks, cotton seed hulls, sawdust, wood chips) has been used as incubation ground and fiber reinforcement for sustainable mycelium-based composite materials for structural applications (e.g. reference ^[31]^ and review by Elsacker et al. ^[32]^). To our knowledge, the use of agricultural waste for promoting the growth of mycelium that could remove antibiotics from contaminated water has not been tested. Thus, the question arises about the synergistic nature of agricultural waste, ambient conditions, and mycelium, not only for structural applications, but also for mycoremediation. To the best of our knowledge, the agricultural waste used in this paper, aged almond shells and fava bean stalks, haw never been used as a mycelium growth substrate for any applications (structural or mycoremediation). The state of California produces 80% of the worldwide almond supply (2016 data), with considerable byproduct in the form of almond shells and hulls for every kg of almond produced. This waste is typically used for livestock feed and biofuel production. Fava beans are a staple food in many countries, and are a common cover crop in USA, replenishing the soil with nitrogen and nutrients. One of the additional objectives of this study was to incorporate biomass sources from local vegetable gardens, as this would serve as a more practical approach to grow biomass on food waste generated locally (versus in specialized industries).

Biomass ingredients may allow the mycelium of a common edible and medicinal fungus, *Ganoderma lucidum*, to thrive. *Ganoderma lucidum* is rarely found in nature, and it is mostly cultivated artificially because it is in high demand for its medicinal properties. ^[33]^ It was chosen in this experiment for a number of reasons: 1) it is mentioned in several works on structural mycelium composites ^[32]^; 2) it has been successfully grown on select agricultural waste biomass; and 3) prior published work on mycoremediation ^[1, 17–18, 22, 30]^ demonstrated its ability to oxidize antibiotics in vitro at various success levels, via extracellular peroxidases, pectinases, ligninase, xylanases, cellulases, and oxidases. We therefore hypothesized that it could be used with the appropriate biomass substrate to remove antibiotics from water.

The mycoremediation experiment of this paper investigates the degradation of twenty antibiotics from the six following classes: Amphenicols, Sulfonamides, B-lactans, Lincosamides, Quinolones, and Macrolides. These antibiotics are commonly detected in wastewater and aquaculture farming (**Table 1**). Quinolones are not biodegradable and remain active in sediments for prolonged periods of time. ^[11, 14, 34]^ Their use has been completely unrestricted in growing aquaculture industries such as China and Chile, where annually 10-12 metric tons of quinolones are used in human medicine, and approximately 100-110 metric tons are used in veterinary medicine per year. ^[34]^

**Table 1.** Antibiotic standards (Emami and Taha ^[35]^)

| *Class* | *Antibiotic Abbreviation* | *Antibiotic Name* | *Catalogue Number* |
| --- | --- | --- | --- |
| **Methanol Soluble:** |  |  |  |
| **Amphenicols** | CAP | *Chloramphenicol* | 40524 |
|  | TAP | *Thiamphenicol* | AAJ63575-03 |
|  | FF | *Florfenicol* | 73231-34-2 |
|  | FFA | *Florfenicol amine* | F405773 |
| **Sulfonamides** | SDM | *Sulfadimethoxine* | 122-11-2 |
|  | SSZ | *Sulfasalazine* | 15025 |
|  | SMX | *Sulfamethoxazole* | 23613 |
|  | SDZ | *Sulfadiazine* | 68-35-9 |
| **Lincosamides** | LIN | *Lincomycin* | 21526 |
| **Quinolones** | ENRO | *Enrofloxacin* | 33699 |
|  | FLU | *Flumequine* | 21645 |
|  | NOR | *Norfloxacin* | 70458-96-7 |
|  | ENO | *Enoxacin* | 16956 |
| **Macrolides** | ERYTH | *Erythromycin* | 114-07-8 |
|  | VIRG-M1 + VIRG-S1 | *Virginiamycin M1 + Virginiamycin S1* | 14503 |
| **Water Soluble:** |  |  |  |
| **B-lactams** | AMP | *Ampicillin* | 69-53-4 |
|  | PEN-G | *Penicillin G* | 21615 |
|  | PEN-V | *Penicillin V* | 23635 |
|  | AMOX | *Amoxicillin* | 26787-78-0 |

## 2. Materials and methods

### 2.1 Materials

LC/MS grade methanol, optima grade methanol, and toluene were obtained from Fisher Chemical, LC/MS grade. Acetonitrile was obtained from Thermo Scientific. Formic acid, HCl 37% and Na_2_EDTA were purchased from Sigma-Aldrich (St. Louis, MO). Dimethyldichlorosilane (DMDCS) was obtained from Fisher Scientific (Hampton, NH, USA).

Antibiotic standards used in this study were from the following classes: Amphenicols (chloramphenicol (CAP), thiamphenicol (TAP), florfenicol (FF), florfenicol amine (FFA)); Sulfonamides (sulfadimethoxine (SDM), sulfasalazine (SSZ), sulfamethoxazole (SMX), sulfadiazine (SDZ)); B-lactams (ampicillin anhydrous (AMP), penicillin G potassium salt (PEN-G), penicillin V (PEN-V) and amoxicillin (AMOX)); Lincosamides (lincomycin (LIN)); Quinolones (enrofloxacin (ENRO), flumequine (FLU), norfloxacin (NOR), enoxacin (ENO)); Macrolides (erythromycin (ERYTH), virginiamycin complex (VIRG M1 and VIRG S1)).

CAP (98.5%) was purchased from Crescent Chemical (Islandia, NY). TAP (99.3%), FF (98%), SDZ (99%) and AMOX (98%) were purchased from Fisher Scientific (Hampton, NH, USA). ERYTH (94.8%), ENRO (99.8%), FFA (99.3%), SDM (98.5%), AMP (99.6%), and NOR (98%) were purchased from Sigma Aldrich (St. Louis, MO). SSZ (100%), SMX (100%), LIN (98%), PEN-V (98.8%), PEN-G, FLU, ENO, and VIRG were purchased from Cayman Chemicals (Ann Arbor, MI).

Isotopically labeled surrogate standards including florfenicol amine-D3 (FFA-D3) (chemical purity: 98%; isotopic purity: 98.7%), chloramphenicol-D5 (CAP-D5) (chemical purity: 98%; isotopic purity: 98.3%), lincomycin-D3 (LIN-D3) (chemical purity: 95%; isotopic purity: 99.6%), sulfamethoxazole-D4 (SMX-D4) (chemical purity: 98%; isotopic purity: 99.2%), sulfamethoxazole-D4 (SMZ-D4) (chemical purity: 98%; isotopic purity: 95.9%), erythromycin (ERYTH-D6) (chemical purity: 95%; isotopic purity: 98.1%) and enrofloxacin-D5 (ENRO-D5) (chemical purity: 99.61%; isotopic purity: 99.40%), ampicillin-D5 (AMP-D5) (chemical purity: 95%; isotopic purity: 99.00%), were purchased from Toronto Research Chemicals (Toronto, Ontario, Canada).

### 2.2 Biomass preparation

The biomass was prepared by mixing 200 mL of oak pellets (MushroomMediaOnline, IA, USA), 200 mL of almond shells from California (leftovers from a research project completed before Fall 2020) and 400 mL of filtered tap water. Used coffee grounds (3 tablespoons) from a local household and 100 mL of fava stalks from a local vegetable garden (a Spring 2021 cover crop harvested, dried in the sun and then pulverized) were added as nitrogen-rich nutrient source. The biomass mix was autoclaved in an autoclaveable bag with a filter patch (grow.bio, USA), at 121 deg. C and at 0.103 MPa (15 psi) for 20 minutes, and then cooled for 1 hour in a fume hood. The autoclaved biomass mix was then inoculated with 5 mL *Ganoderma lucidum* liquid culture (Root Mushroom Farm, WA, USA). After 13 days in ambient conditions (October 2021 in Davis, CA), sustained growth of the mycelium into the biomass was observed (**Figure 1**), and the bag with mycelium and biomass was relocated to a −80 deg. C freezer for storage.

**Figure 1.**
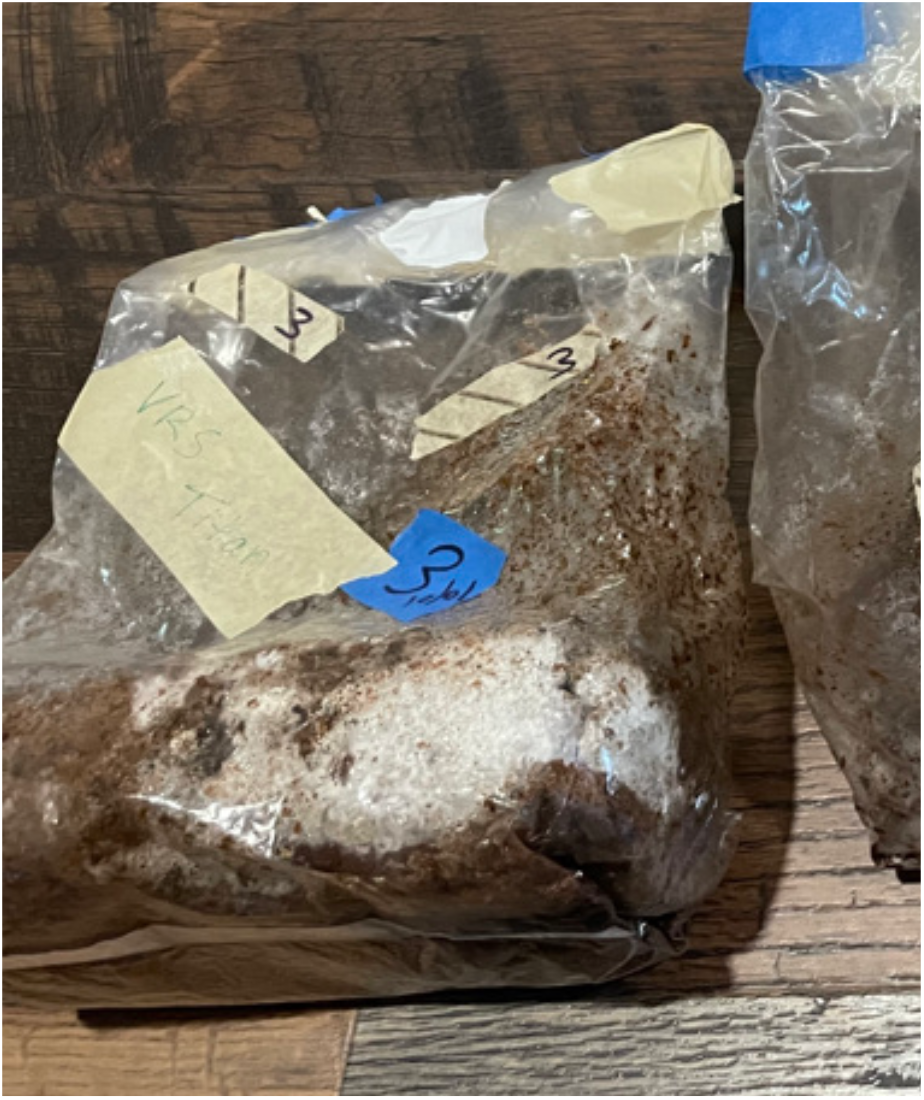
Mycelial biomass used in this study.

### 2.3 Antibiotic mixture preparation

Antibiotics were first prepared individually in methanol (amphenicols, quinolones, sulfonamides and macrolides) or water (B-lactams). Individual stock solutions of CAP, TAP, FF, FFA, SDM, SMX, ENRO, ERYTH, VIRG, LIN, CAP-D5, SMX-D4, SMZ-D4, ERYTH-D6, ENRO-D5 and TRIM-D3 were prepared in methanol at 1 mg/mL concentration. SSZ, SDZ, and LIN-D3 were prepared in methanol at a concentration of 0.5 mg/mL. ENO, NOR and FLU were prepared in methanol at concentration of 0.2 mg/mL. Β-lactams (AMP, PEN-G, PEN-V, AMOX and PEN-V-D5) were prepared in Milli-Q water at 1 mg/mL. AMP-D5 was prepared in Milli-Q water at 0.5 mg/mL. The stock solutions were diluted from 0.2 - 1 mg/mL to individual ‘intermediate’ solutions of 10 μg/mL using the same solvent as the stock solution.

The individual intermediate solutions of unlabeled and labeled standards were used to prepare antibiotic mixture solutions of methanol-soluble and water-soluble antibiotic standards. These mixes were prepared separately for unlabeled and labeled antibiotics. For unlabeled antibiotics, both methanol-soluble and water-soluble mixes were prepared at concentration of 400 ng/mL. For labeled antibiotics, both methanol-soluble and water-soluble mixes were prepared at concentration of 1000 ng/mL. The water-soluble and methanol-soluble antibiotics were mixed at a 1:1 ratio, before use. Note that unlabeled and labeled standards were mixed separately, as they were needed for different purposes: the unlabeled mixes were used for spiking water samples, and the labeled mixes were spiked to each sample right before extraction, for the purpose of quantification.

Briefly, for methanol-soluble unlabeled antibiotic standards (n=15; **Table 1**), 30 µL of each antibiotic from their individual intermediate solutions (10 µg/mL) were added to a 2 mL amber glass LC (liquid chromatography) vial. The mixture was evaporated under nitrogen and reconstituted in 750 µL LC-MS ((liquid chromatography-mass spectrometry) methanol. For water-soluble unlabeled antibiotic standards (n=4; **Table 1**), 630 µL Milli-Q water was added to a 2 mL amber glass LC vial followed by adding, 30 µL of each of the four unlabeled water-soluble antibiotic standards (from their individual intermediate solutions (10 µg/mL).

For methanol-soluble labeled antibiotic standards (n=7; **Table 2**), 50 µL of each labeled standard (from their individual intermediate solutions (10 µg/mL) were added to a 2 mL amber glass LC vial. Samples were vortexed, dried under nitrogen, and reconstituted in 500 µL LC-MS methanol. For water-soluble labeled antibiotic standard (n=2; **Table 2**), 400 µL Milli-Q water was added to a 2 mL amber glass LC vial followed by adding 50 µL of the two water soluble standards from their individual intermediate solutions (10 µg/mL). Water-soluble and methanol-soluble antibiotic mixes were mixed at a 1:1 ratio before experiment, resulting in working mix of unlabeled antibiotic standards at concentration of 200 ng/mL and working mix labeled standards at concentration of 500 ng/mL.

**Table 2.** Surrogate standards (Emami and Taha ^[35]^)

| Surrogate Abbreviation | Surrogate Name | Catalogue Number |
| --- | --- | --- |
| *Methanol Soluble:* |  |  |
| **CAP – D5** | *Chloramphenicol – D5* | C325033 |
| **FFA – D3** | *Florfenicol amine – D3* | F405773 |
| **TRIM – D3** | *Trimethoprim – D3* | T79618 |
| **SMZ – D4** | *Sulfamethazine – D4* | S699072 |
| **SMX – D4** | *Sulfamethoxazole – D4* | S699087 |
| **LIN – D3** | *Lincomycin – D3* | L466202 |
| **ENRO – D5** | *Enrofloxacin – D5* | E557802 |
| **ERYTH – D6** | *Erythromycin – D6* | E649953 |
| *Water Soluble:* |  |  |
| **AMP – D5** | *Ampicillin – D5* | A634337 |
| **PEN-V – D5** | *Penicillin V – D5* | 26786 |

### 2.4 Glass silanization prior to antibiotic treatment

Some antibiotics including quinolones were shown to adsorb onto the glass surface, likely by interactions with the glass silanol groups. ^[35]^ In order to avoid these interactions, glassware was silanized using dimethyldichlorosilane (DMDCS) prior to the experiment, in order to cover the silanol groups on the glass surface.

Glassware were silanized using the method described by Ye et al. ^[36]^. Briefly, the clean and dry glass vials (50 mL), were pre-washed with detergent-free soap and water and rinsed 5 times with distilled water, were treated with 5 mL 5% DMDCS in toluene solution. Silanization was completed by rinsing, vortexing, and dumping three more times with 5 mL of 5% DMDCS in toluene, 5 mL toluene, 5 mL methanol, and 5 mL Milli-Q water; respectively. Vials were left overnight to dry.

### 2.5 Experimental design and mycelium treatment

Mycelial biomass (*Ganoderma lucidum* grown on the substrate described in Section 2.2) was thawed on ice for approximately 90 minutes. Approximately, 1 g of biomass was weighed and placed into 50 mL silanized Pyrex glass tubes. Milli-Q water (50 mL) was then added. Mycelium-water samples were spiked with 20 ng per 25 mL water of unlabeled antibiotics (**Table 1**) and incubated for 0 (baseline) or 3 days (n=4 samples per incubation period). A parallel set of samples (n=4 samples per timepoint) contained 20 ng antibiotics in 25 mL Milli-Q water without mycelium, as a negative control. Each sample was set in its own independent 50 mL Pyrex tube. Sample corresponding to Day 3 were incubated for 3 days, then on Day 3, the Day 0 (baseline) samples was completed (in separate Pyrex tubes). Additionally, one water method blank containing 25 mL of Milli-Q water and no antibiotics (n=1 on Day 0 and Day 3), as well as one matrix blank containing mycelium and water only (no antibiotics; n=1 on Day 0 and n=1 on Day 3) were incorporated into the design to control for any background antibiotics potentially coming from the water itself or the mycelium, respectively. Antibiotics were extracted from both Day 0 and Day 3 samples as described below. Vials representing “Day 3” sample were covered in foil and shaken at room temperature using an Excella E24 shaker at 100 rpm for 3 days.

The final design and sample size for each incubation day (i.e. Day 0 baseline, and Day 3) were as follows:

n= 1, Water method blank (Water only) = 25 mL Milli-Q water

n= 1, Matrix method blank (Water + Mycelium, or “WM” group) = 0.5 g mycelial biomass in 25 mL Milli-Q water

n= 4, Control sample (Water + Antibiotics, or “WA” group) = 20 ng antibiotics in 25 mL Milli-Q water

n= 4, Treatment sample (Water + mycelial biomass + Antibiotics, or “WMA” group) = 0.5 g mycelial biomass + 20 ng antibiotics in 25 mL Milli-Q water

After incubation, the samples were centrifuged using SORVALL RT 6000D at 3000 rpm for 10 minutes at room temperature. One water-mycelial biomass Day 3 (WM-D3) sample and one water-mycelial biomass antibiotics Day 3 (WMA-D3) sample broke in the centrifuge, and the WM-D3 sample was unrecoverable. To protect the rest of the samples, including all those from Day 0, samples were just vortexed. Water samples were then transferred to 40 mL non-silanized glass vials for the extraction of antibiotics.

### 2.6 Antibiotics extraction

25 mL water samples placed in non-silanized glass vials were spiked with 20 ng of surrogate standard mixture containing CAP-D5, FFA-D3, TRIM-D3, SMZ-D4, SMX-D4, LIN-D3, ENRO-D5, ERYTH-D6, AMP-D5, and PEN-V-D5 (500 ng/mL per surrogate standard, **Table 2**). Then, 1340 µL of 0.05 M Na_2_EDTA in water (corresponding to 0.1% Na_2_EDTA) was added to glass vials. Aliquots were let to stay for 1 hour covered with aluminum foil with occasional shaking before solid phase extraction (SPE).

### 2.7 Solid Phase Extraction (SPE) and sample preparation

Antibiotics were extracted from the water samples using Waters OASIS HLB cartridges (60 mg, 3 cm). The cartridges were pre-conditioned using methanol (5mL), Milli-Q water (5mL), and pH = 2.5 water (5mL) made by adding 90 µL HCL (37%) to 150 mL Milli-Q water in a flask, and verifying the pH with litmus paper. Then, water samples were loaded onto the conditioned HLB cartridges and allowed to elute. The SPE cartridges were washed with 6 mL Milli-Q water. The cartridges were then dried under the vacuum manifold (Supelco Visiprep 24 SPE) for five minutes at 0.117 MPa (17 psi). Antibiotics were eluted into 8 mL glass vials using 5 mL optima grade methanol. The vials were stored overnight in −20 deg. C freezer. The following day, the samples were evaporated under nitrogen for two hours. Samples were reconstituted in 1 mL LC-MS methanol: water (1:1). All samples (n=17) from the SPE step were vortexed for 3 minutes and then transferred to 2 mL centrifuge tubes. The tubes were centrifuged for 2 min at 12000 rpm at 0 deg. C (Eppendorf, 5424 R, 13523 ×g). The samples were then transferred into filter centrifuge tubes (two tubes per sample each containing (approx. <500 µL of sample each tube). The actual sample amount pipetted was 480 µL per tube. The filter centrifuge tubes containing the samples were centrifuged for 10 minutes at 12000 rpm (Eppendorf, 5424 R, 13523 ×g). After centrifugation, the filters were discarded, and the extract was transferred to inserts containing LC vials with slit caps to be analyzed by UPLC-MS/MS (Ultra performance liquid chromatography with tandem mass spectrometry).

### 2.8 LC-MS/MS instrumentation

Antibiotic analysis was performed on an Agilent ultra-high pressure liquid chromatography coupled to a 6460 Agilent triple quad (LC-MS/MS). Chromatographic separation of the antibiotics mixture was performed on AQUITY BEH C18 column (100 × 2.1 mm, 1.8 µm), using 0.1% formic acid in water (mobile phase A, (MPA)) and 0.1% formic acid in acetonitrile (mobile phase B, (MPB)) running at flow rate of 0.300 mL/min and column temperature of 30 deg. C. MS/MS (tandem mass spectrometry) analysis was performed using Agilent Jetstream electrospray ionization (ESI) operating at both positive and negative mode. MS source parameters were as follows: sheath gas temperature of 375 deg. C, sheath gas flow of 8 L/min, drying gas temperature of 250 deg. C, nozzle voltage of 0 V, nebulizer gas pressure of 40 psi and capillary voltage of 3500 V (**Table 4S.** Source Parameters for LC/MS (Emami and Taha ^[35]^)). Collision-induced dissociation was carried out using nitrogen at the collision cell.

### 2.9 Antibiotics quantification

Antibiotic concentrations in water samples were calculated by the internal standard calibration method where isotopically labeled surrogates were used to correct for both recoveries and the detector response factor. A 11-point standard calibration curve (0.01-100 ng/mL) containing a fixed amount of surrogate standard (20 ng/mL) was made to derive the response factor. Calibration curves were generated by quadratic regression and a 1/x^2^ weighting factor was applied. Peaks were integrated and analyzed using MassHunter Workstation Software, QQQ Quantitative Analysis (Agilent Technologies, Inc.).

### 2.10 Matrix effects

Matrix effects (ME) from water-mycelium matrix were assessed for the 20 selected antibiotics at Day 0 (D0) spiked at a concentration of 20 ng per 25 mL of water (ME-D0, n=2). Matrix effects were calculated as the ratio of the peak area of the antibiotic standards (unlabeled and labeled) in sample spiked after extraction (ME-D0) to the peak area of antibiotic standards in pure solvent (**Table 3**).

**Table 3.** Matrix effects for 20 antibiotic standards (unlabeled standard) and 10 isotopically labeled standards in Day 0. Matrix effects were calculated by dividing the peak area (response) in Water-Mycelium samples (WM) spiked with standards right after extraction (ME, n=2) to the peak area in pure solvent.

| Antibiotic Abbreviation | Antibiotic Name | Matrix Effects: Day 0 Sample 1 | Matrix Effects: Day 0 Sample 2 |
| --- | --- | --- | --- |
| **CAP-D5** |  | **99%** | **16%** |
| CAP | *Chloramphenicol* | 121% | 46% |
| TAP | *Thiamphenicol* | 31% | 7% |
| FF | *Florfenicol* | 64% | 20% |
| **FFA-D3** |  | **59%** | **30%** |
| FFA | *Florfenicol amine* | 43% | 26% |
| **SMX-D4** |  | **142%** | **45%** |
| SMX | *Sulfamethoxazole* | 95% | 49% |
| SDM | *Sulfadimethoxine* | 87% | 49% |
| VIRG-M1 | *Virginiamycin M1* | 90% | 64% |
| VIRG-S1 | *Virginiamycin S1* | 158% | 168% |
| **SMZ-D4** |  | **81%** | **22%** |
| SSZ | *Sulfasalazine* | 66% | 47% |
| SDZ | *Sulfadiazine* | 50% | 25% |
| **AMP-D5** |  | **116%** | **73%** |
| AMP | *Ampicillin* | 79% | 55% |
| AMOX | *Amoxicillin* | 83% | 61% |
| **PEN-V-D5** |  | **84%** | **37%** |
| PEN-G | *Penicillin G* | 136% | 77% |
| PEN-V | *Penicillin V* | 108% | 79% |
| **LIN-D3** |  | **132%** | **54%** |
| LIN | *Lincomycin* | 105% | 69% |
| **ENRO-D5** |  | **42%** | **21%** |
| ENRO | *Enrofloxacin* | 48% | 48% |
| NOR | *Norfloxacin* | 126% | 181% |
| ENO | *Enoxacin* | 87% | 108% |
| **TRIM-D3** |  | **97%** | **36%** |
| FLU | *Flumequine* | 104% | 80% |
| **ERYTH-D6** |  | **65%** | **13%** |
| ERYTH | *Erythromycin* | 50% | 16% |

### 2.11 Statistical analysis

Data was analyzed using MassHunter Workstation software, QQQ Quantitative Analysis (Agilent Technologies, Inc.). To compare the recoveries, results were exported to an Excel spreadsheet where percent changes between Day 0 and Day 3 water-mycelial biomass-antibiotics (WMA) and water antibiotics (WA) were collected. The commercial software MATLAB (Mathworks) performed three normality tests (Lilliefors, Anderson-Darling and Kolmogorov-Smirnov), a Mann-Whitney/Wilcoxon test and boxplots for non-parametric distributions.

## 3. Results and discussion

### 3.1 Normality test

The statistical analyses showed that the normality tests had mixed results (Lilliefors and Anderson-Darling tests ruling many of the distributions as normal, Kolmogorov-Smirnov always treating them as not normal, at 95% confidence levels). Data were analyzed by Mann-Whitney/Wilcoxon test. Boxplots were chosen for the presentation of the data, as very visual straightforward non-parametric representations.

**Fig. *2*** shows boxplots for significant changes in median antibiotics between Days 3 and Day 0 for WMA (treated mycelial biomass water in antibiotics) and WA (water in antibiotics). The normalized median changes are (see also **for LC**^/MS^ (Emami and Taha ^[35]^)) −61.9%, −82.4% and −80.8% respectively for three quinolones (enoxacin ENO, enrofloxacin ENRO, norfloxacin NOR), and −37.7%, −24.4% and −39.5% respectively for three sulfonamides (sulfadimethoxine SDM, sulfasalazine SSZ, sulfadiazine SDZ). For the other antibiotics, the results were statistically equivalent (**Figure 2S**. Boxplots for CAP tests).

**Fig. 2.**
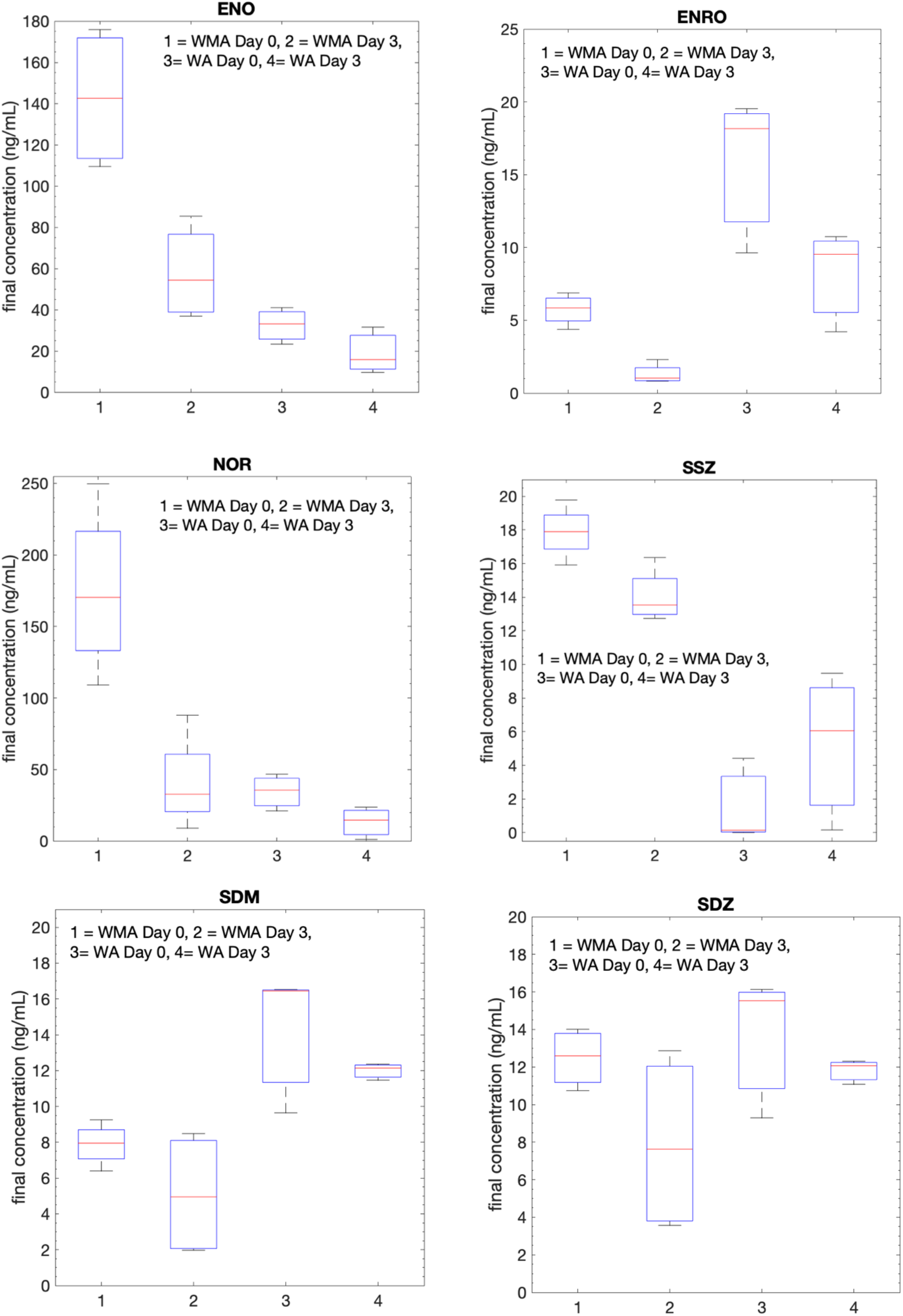
Boxplots of final concentrations of six antibiotics in water treated with mycelial biomass (WMA) or lacking mycelial biomass (WA). Legend: enoxacin ENO, enrofloxacin ENRO, norfloxacin NOR, sulfadimethoxine SDM, sulfasalazine SSZ, sulfadiazine SDZ.

Concentrations for quinolones including NOR and ENO in Day 0 samples (n=4) were higher than the expected concentration of 20 ng/mL (**Fig. *2***). The mean concentration of NOR in Water-Mycelium-Antibiotic (WMA) samples (n=4) was 198.59 ng/mL ± 51.11. The mean concentration of ENO was 161.35 ng/mL ± 48.49. In assessing matrix effects for the three significant quinolones (enrofloxacin ENRO, norfloxacin NOR, and enoxacin ENO), the surrogate ENRO-D5 (enrofloxacin-D5) was used. Analytes NOR and ENO show different matrix effects compared to the surrogate ENRO-D5. Particularly, ENO did not show any matrix effects (87% and 108%), NOR showed ion enhancement (126% and 181%), and ENRO-D5 showed ion suppression (42% and 21%). The discrepancies in the matrix effects of the analytes NOR and ENO (quinolones) and surrogate ENRO-D5 may affect the accuracy in concentrations, resulting in overestimated concentrations.

ENRO did not exhibit high Day 0 concentrations as the other two quinolones. This coincides with ENRO and ENRO-D5 behaving similarly in terms of matrix effects (**Table 3**). Low concentrations found for ENRO (6.14 ng/mL ± 0.62) may be due to quick adsorption to biomass on Day 0. Future studies are required to test whether the biomass is adsorbing the antibiotics and therefore reducing the initial concentration levels.

It should be noted that the matrix effects results (ME 1 and ME 2, **Table 3**) showed high variation. This is likely because these samples were only vortexed and not centrifuged. This could possibly result in inconsistent extraction of matrix components into the supernatant phase, and thereby variable matrix effects.

### 3.2. Results

With respect to published results in mycelium cultures without agricultural waste, Vasiliadou et al. ^[22]^ reported a less than −20% change after an exposure of 7 days to sulfamethoxazole (SMX) by *Ganoderma lucidum* grown with malt extract. It is possible that our results (−32.8% for WMA and −27.8% for WA) may be due to different experimental procedures, or it could be due to the synergistic chemistry of mycelium/agricultural waste mixture. Chakraborty and Abraham ^[30]^ reported 100% removal after 4 days of exposure to enrofloxacin (ENRO) by *Ganoderma lucidum*, findings which are comparable to our results (−82.4% after 3 days in WMA). Martens et al. ^[28]^ showed considerably less success rate in degrading enrofloxacin (ENRO) by their mycelial species (varying between 0.19% for white rot fungus, and 25.6% for brow rot fungus, and after 56 days), likely due to differences in the species used to degrade antibiotics. Finally, Čvančarová et al. ^[21]^ showed that norfloxacin (NOR) degraded by up to 100% with *Irpex lacteus* after 10 days, and by 10% with *Dichomitus squalens* after 10 days. Due to the considerably different experimental conditions, we cannot directly compare the outcome of our work with research by bioreactors based on fungal pellets. ^[24–26, 29]^

Our results suggest that we have identified a promising mycoremediation process that warrants further investigation. Clearly, different fungi strains degrade antibiotics with different capacities.

## 4. Conclusions

In our investigation, ultra-high pressure liquid chromatography coupled to tandem mass-spectrometry determined whether a mycelium (*Ganoderma lucidum*) grown on agricultural waste could impact the concentrations of twenty common antibiotics in synthetic wastewater. In just 3 days, the concentration was reduced in six antibiotics (three quinolones and three sulfonamides) with respect to water with no mycelium. Results showed normalized median changes of −61.9%, −82.4% and −80.8% respectively for three quinolones (enoxacin ENO, enrofloxacin ENRO, norfloxacin NOR), and −37.7%, −24.4% and −39.5% respectively, for three sulfonamides (sulfadimethoxine SDM, sulfasalazine SSZ, sulfadiazine SDZ). The other 14 antibiotics showed insignificant degradation, or even increased antibiotic concentrations. The results on quinolone antibiotics are particularly promising because of their extensive consumption worldwide, and the inability of conventional wastewater treatment plants to remove them, thus fostering antibiotics resistance and harming the environment.

Matrix effects show that the ionization of the antibiotic analytes is affected by matrix compounds within the sample extracts due to lack of silanization of Day 0 samples. Stability is known to be affected by temperature, solvent composition and, in this case, container type. ^[37]^ However, there were good extraction recoveries observed for the majority of compounds. Lower recoveries could be caused by the presence of organic matter in the matrix, with surfactant properties that could increase signal intensity, and cause suppression by promoting ionization in the positive electrospray. ^[38]^ Higher recoveries are found to possibly be due to more antibiotic adsorption into the biomass than the mycelium.

Our findings suggest that further studies should be carried out to test for bacterial pollution, since the biomass partly originates from products which have bacteria that could potentially degrade the antibiotics as well, even though the biomass was sterilized before inoculation. In future work, we will use a separate control of water-uninoculated biomass-antibiotics to test whether the antibiotic pollutants are adsorbing into the biomass. Also, our future studies will use a control of water + inactive mycelium (de-activated by heat) + antibiotics, to test if there is biodegradation or only elimination of the liquid medium. Colonized agar plates with potato dextrose will also be used in our next experimental trials as other studies have used in their experiments.

Our work indicates that mycelium and re-purposed agricultural waste is a promising, novel method to remove antibiotics from water.

## Abbreviations

AMOX: Amoxicillin
AMP: Ampicillin
CAP: Chloramphenicol
ENRO: Enrofloxacin
ENO: Enoxacin
ERYTH: Erythromycin
FF: Florfenicol
FFA: Florfenicol amine
FLU: Flumequine
LIN: Lincomycin
NOR: Norfloxacin
PEN-G: Penicillin G
PEN-V: Penicillin V
SPE: Solid phase extraction
SDM: Sulfadimethoxine
SDZ: Sulfadiazine
SMX: Sulfamethoxazole
SSZ: Sulfasalazine
TAP: Thiamphenicol
VIRG-M1: Tilmicosin: Virginiamycin M1
VIRG-S1: Virginiamycin S1

## Author contributions

V.S., V.L.S., S.E. and A.Y.T designed the experiments. V.S. and S.E. performed the experiments and analyzed the data. V.S. wrote the original draft. V.S. and S.E. edited the manuscript. V.L.S. and A.Y.T. reviewed and edited the manuscript and carried out the statistical analysis.

## Data availability statement

The data is available within the article and its supplementary material.

## Supporting materials

We provide additional information in **Tables 1S-5S, Figures 1S-15S**.

### LC-MS/MS run

**Table 1S.** Unlabeled standard and labeled surrogate mix concentrations (Emami and Taha ^[35]^).

| Calibration Level | Methanol:Water (1:1) | Unlabeled - Standard Mix | Labeled - Surrogate Mix |
| --- | --- | --- | --- |
| 0.001 ng/mL | 1919 µL | 250 µL | 80 µL |
| 0.01 ng/mL | 1910 µL | 100 µL | 80 µL |
| 0.1 ng/mL | 1820 µL | 10 µL | 80 µL |
| 0.25 ng/mL | 1670 µL | 1 µL | 80 µL |
| 0.5 ng/mL | 1915 µL | 5 µL | 80 µL |
| 1 ng/mL | 1910 µL | 10 µL | 80 µL |
| 2 ng/mL | 950 µL | 10 µL | 40 µL |
| 5 ng/mL | 374 µL | 10 µL | 16 µL |
| 10 ng/mL | 364 µL | 20 µL | 16 µL |
| 20 ng/mL | 344 µL | 40 µL | 16 µL |
| 50 ng/mL | 284 µL | 100 µL | 16 µL |
| 100 ng/mL | 184 µL | 200 µL | 16 µL |

**Table 2S.** LC-MS/MS order run

| ***Plate Name*** | ***Sample Name*** |
| --- | --- |
| P1A1 | *Blank1.d* |
| P1B2 | *Std50inj1.d* |
| P1B2 | *Std50inj2.d* |
| P1A1 | *Blank2.d* |
| P1A2 | *Std0.01.d* |
| P1A3 | *Std0.1.d* |
| P1A4 | *Std0.25.d* |
| P1A5 | *Std0.5.d* |
| P1A6 | *Std1.d* |
| P1A7 | *Std2.d* |
| P1A8 | *Std5.d* |
| P1A9 | *Std10.d* |
| P1B1 | *Std20.d* |
| P1B2 | *Std50.d* |
| P1B3 | *Std100.d* |
| P1A1 | *Blank3.d* |
| P1A1 | *Blank4.d* |
| P1B4 | *WM-D0.d* |
| P1B5 | *WMA-D0-1.d* |
| P1B6 | *WMA-D0-2.d* |
| P1B7 | *WMA-D0-3.d* |
| P1B8 | *WMA-D0-4.d* |
| P1B9 | *WMA-D3-1.d* |
| P1C1 | *WMA-D3-2.d* |
| P1C2 | *WMA-D3-3.d* |
| P1C3 | *WMA-D3-4.d* |
| P1A1 | *Blank5.d* |
| P1C4 | *WA-D0-1.d* |
| P1C5 | *WA-D0-2.d* |
| P1C6 | *WA-D0-3.d* |
| P1C7 | *WA-D3-1.d* |
| P1C8 | *WA-D3-2.d* |
| P1C9 | *WA-D3-3.d* |
| P1D1 | *ME1.d* |
| P1D2 | *ME2.d* |
| P1A1 | *Blank6.d* |

**Table 3S.** Mobile phases (Emami and Taha ^[35]^) A: 0.1 % formic acid in water B: 0.1 % formic acid in Acetonitrile

|  | **Time** | **A** | **B** | **Flow** | **Pressure** |
| --- | --- | --- | --- | --- | --- |
| 1 | 8.00 min | 80.00 % | 20.00 % | 0.300 mL/min | 600.00 bar |
| 2 | 11.00 min | 60.00 % | 40.00 % | 0.300 mL/min | 600.00 bar |
| 3 | 13.00 min | 0.00 % | 100.00 % | 0.300 mL/min | 600.00 bar |
| 4 | 15.00 min | 0.00 % | 100.00 % | 0.300 mL/min | 600.00 bar |
| 5 | 17.00 min | 90.00 % | 10.00 % | 0.300 mL/min | 600.00 bar |
| 6 | 20.00 min | 90.00 % | 10.00 % | 0.300 mL/min | 600.00 bar |

**Table 4S.** Source Parameters for LC/MS (Emami and Taha ^[35]^)

| **Parameter** | **Value (+)** | **Value (-)** |
| --- | --- | --- |
| Gas Temp (°C) | 250 | 250 |
| Gas Flow (l/min) | 8 | 8 |
| Nebulizer (psi) | 40 | 40 |
| SheathGasHeater | 375 | 375 |
| SheathGasFlow | 11 | 11 |
| Capillary (V) | 3500 | 3500 |
| Vcharging | 0 | 0 |

**Table 5S.**
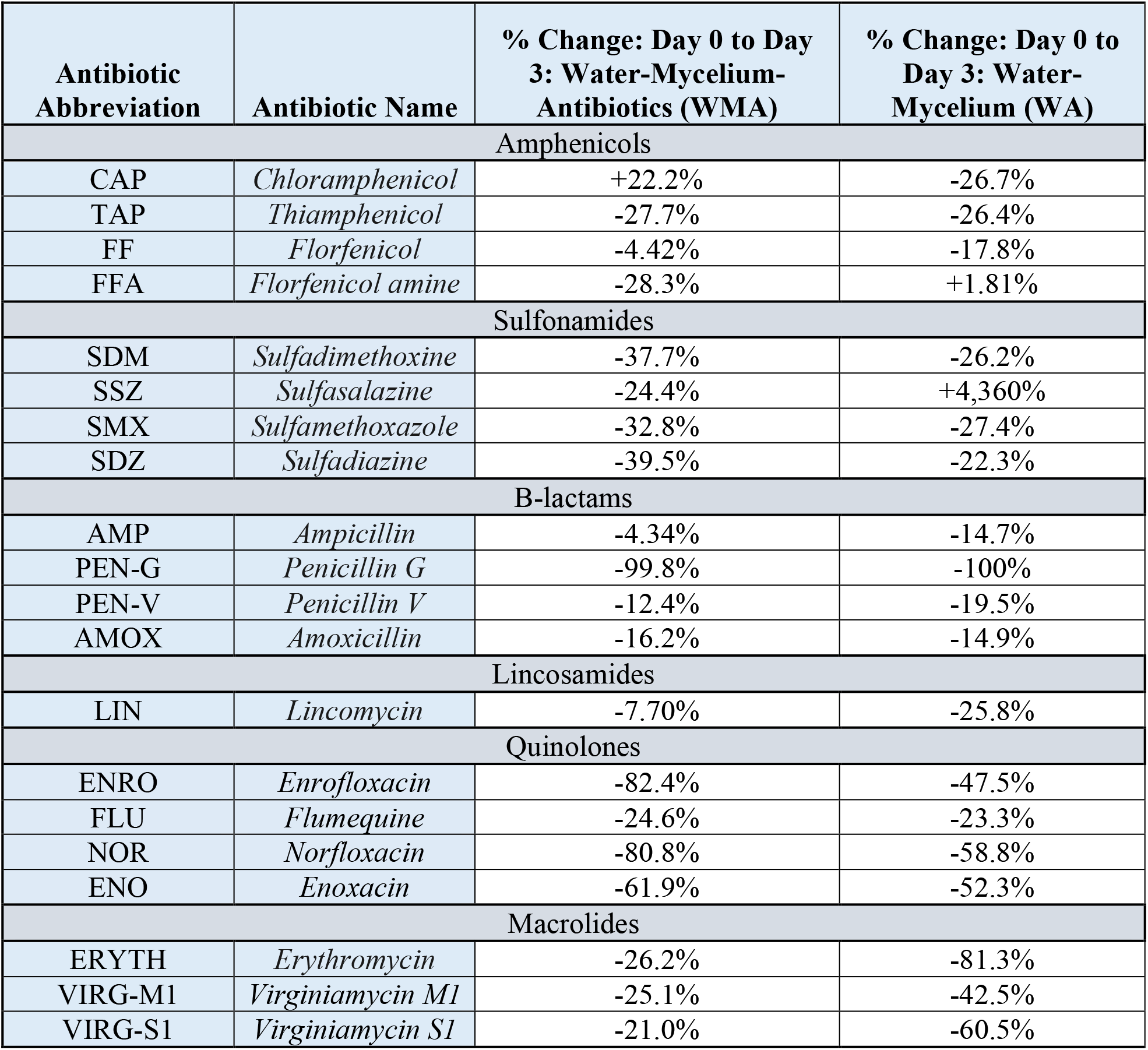
Percent normalized changes between day 0 to day 3 medians of Water-Mycelium-Antibiotics (WMA) and Water-Mycelium (WA) groups.

**Figure 1S.**
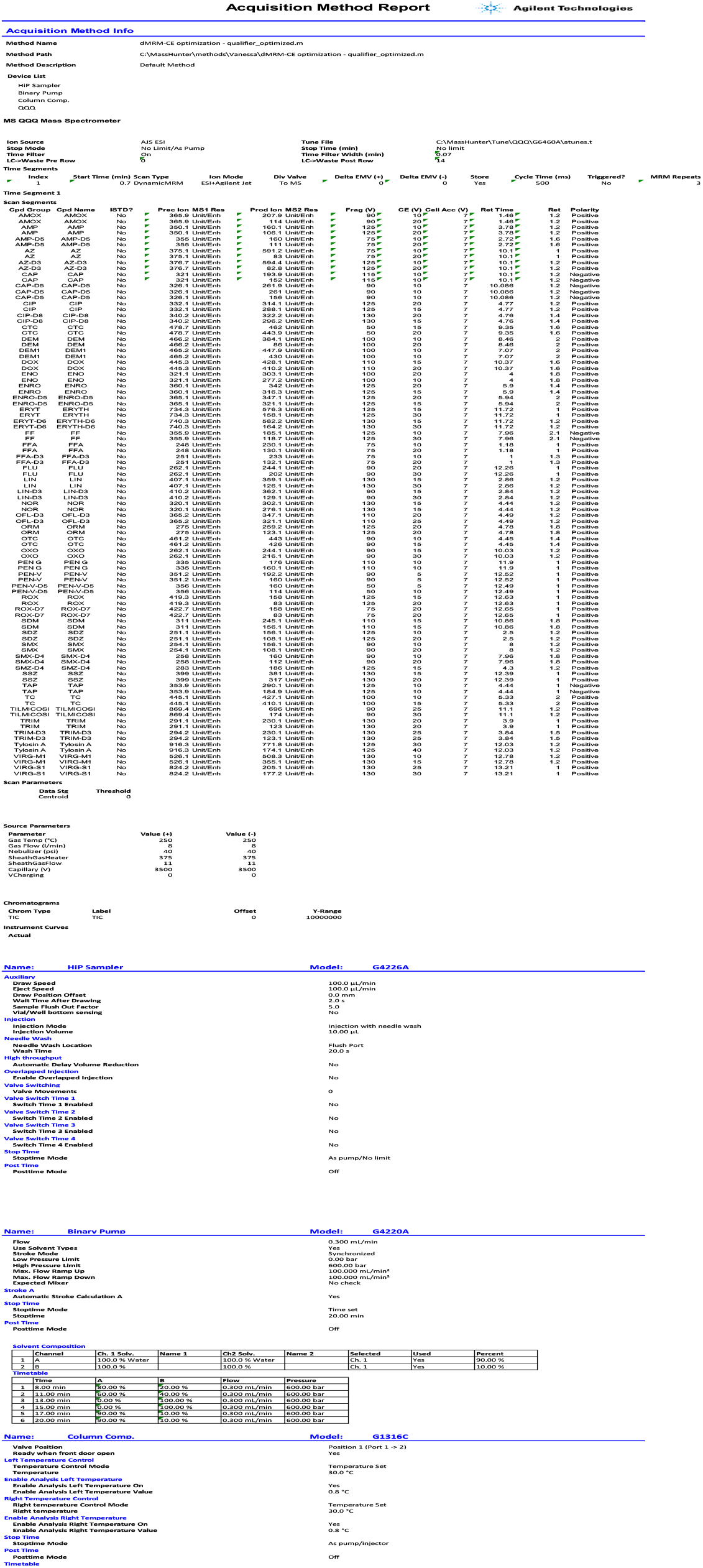
Acquisition Method Report. The boxplots for the final concentrations (in ng/mL) of 14 antibiotics are shown in **Figures 2S-15S** (all except for quinolones ENO, ENRO, NOR, and sulfonamides SSZ, SDM, SDZ, which are shown in **Fig. *2***.) WMA is the synthetic wastewater treated with mycelial biomass, WA contains no mycelial biomass.

**Figure 2S.**
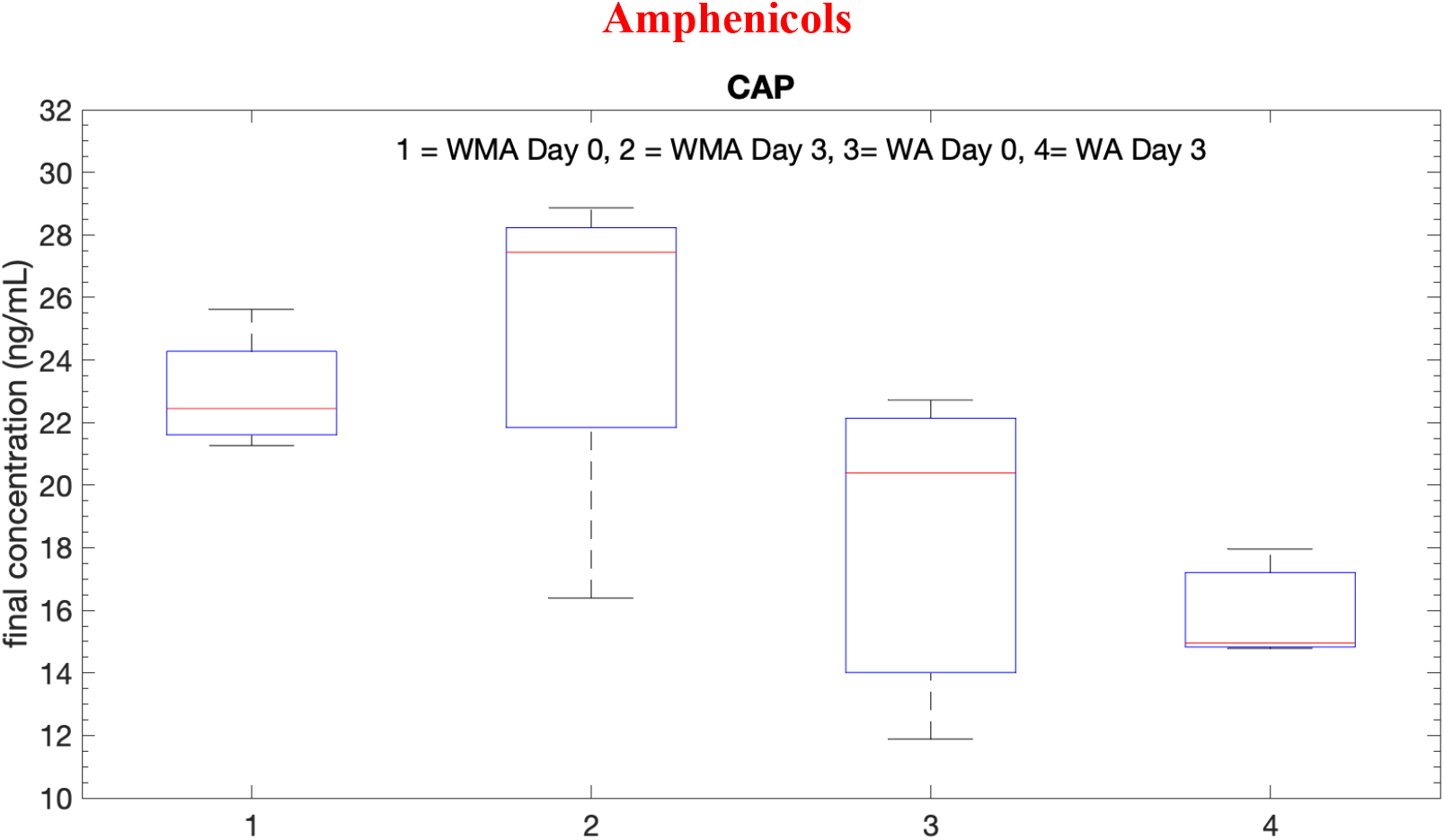
Boxplots for CAP tests

**Figure 3S.**
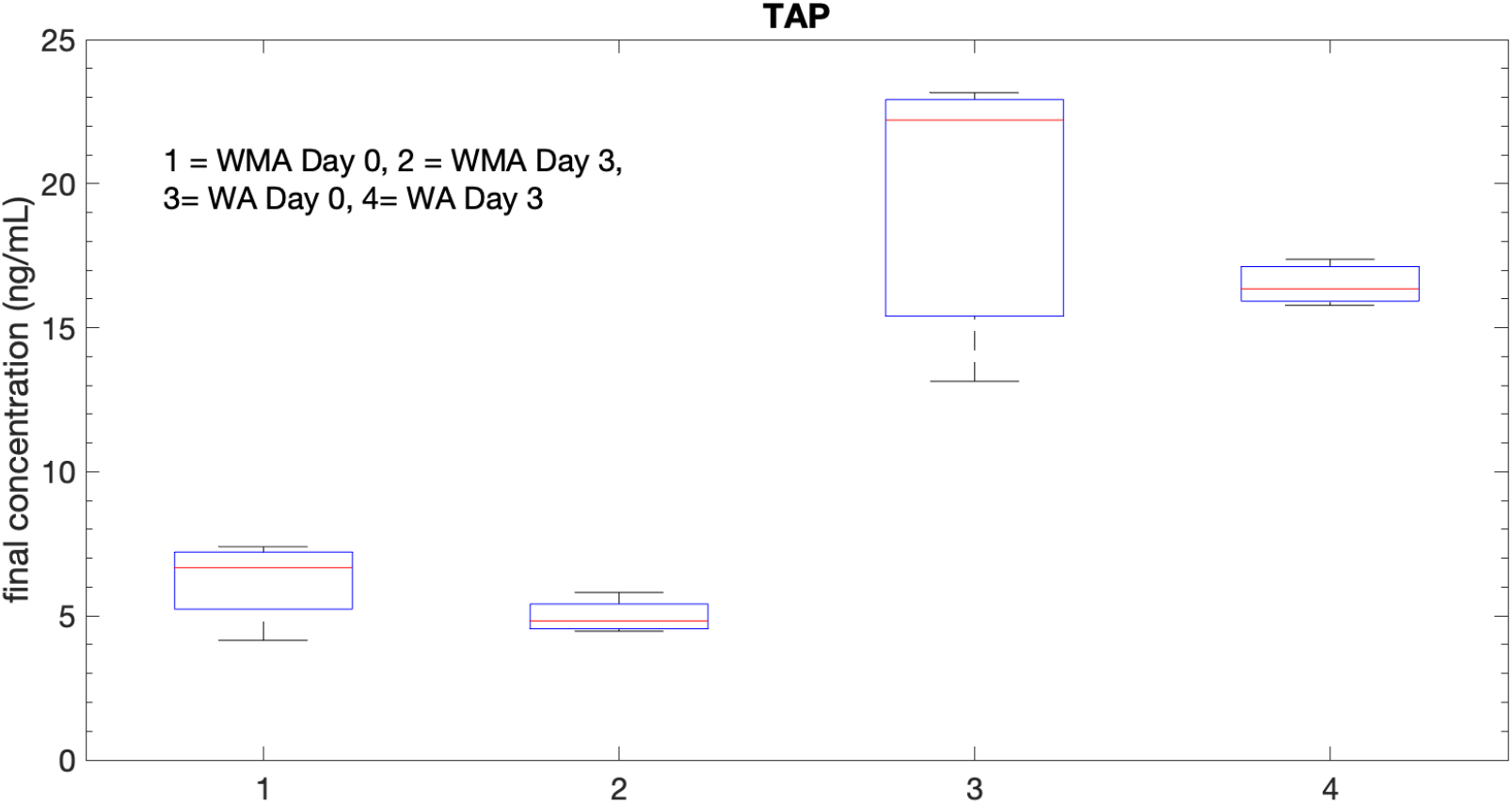
Boxplots for TAP tests.

**Figure 4S.**
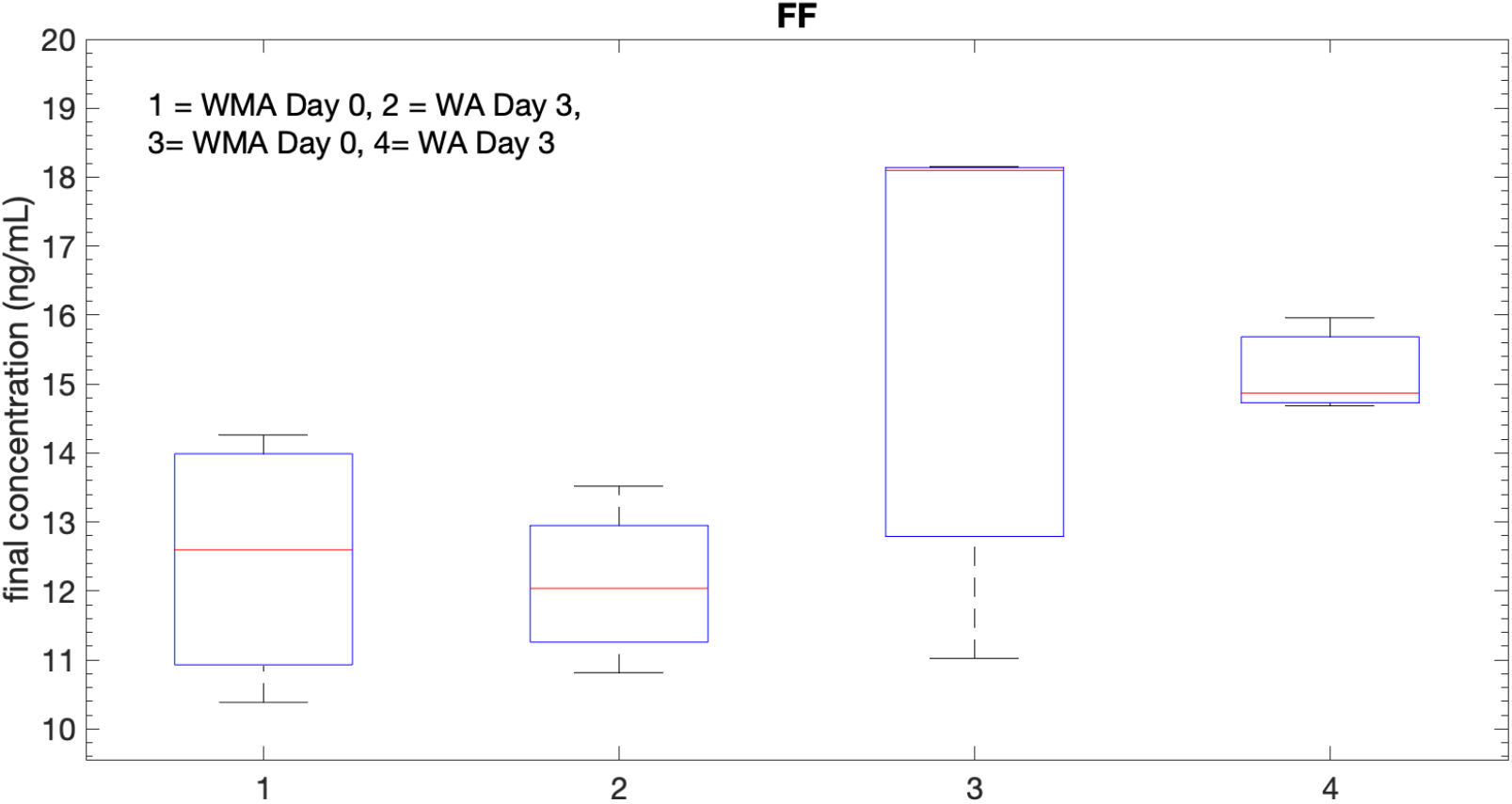
Boxplots for FF tests.

**Figure 5S.**
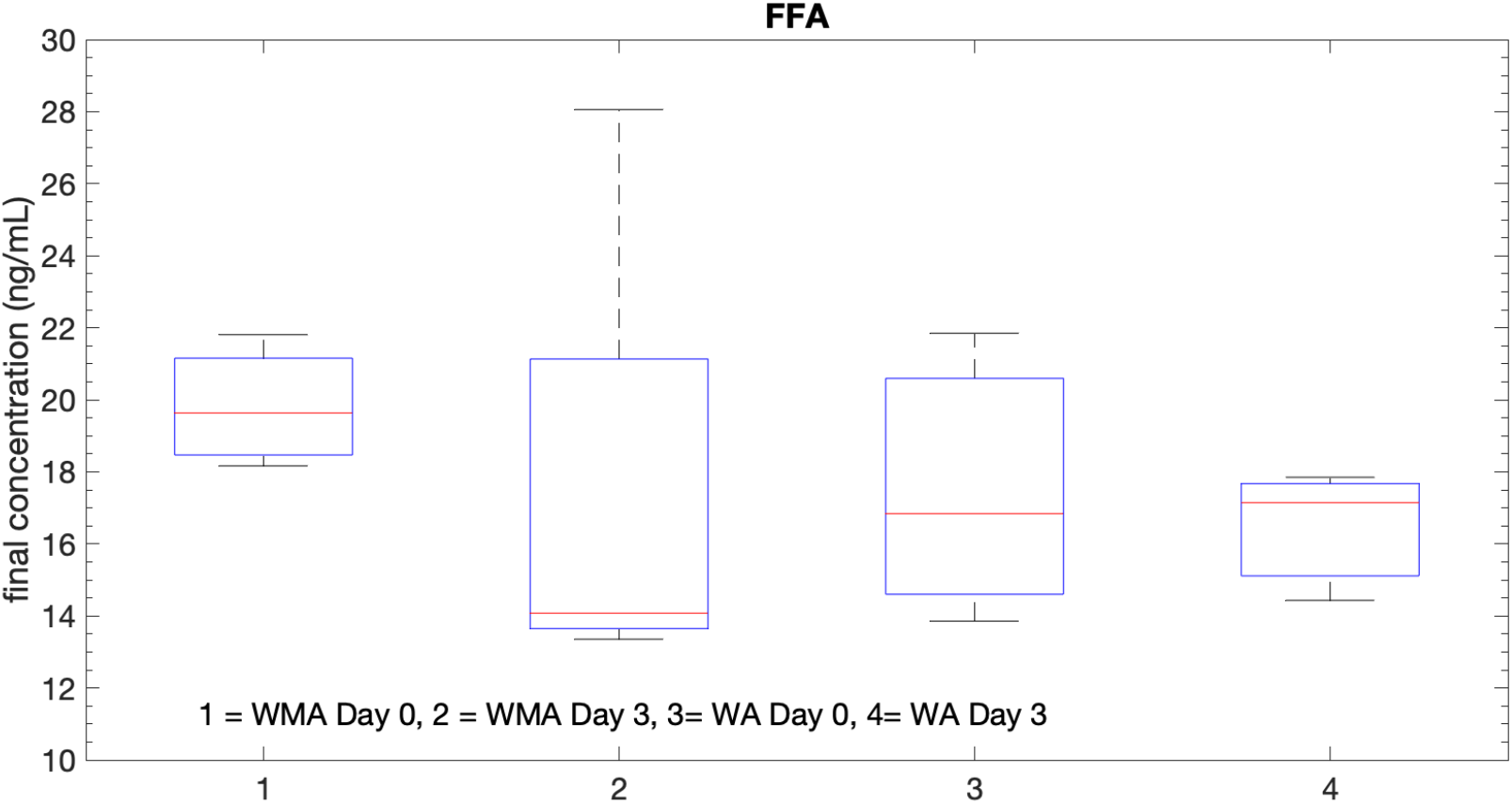
Boxplots for FFA tests.

**Figure 6S.**
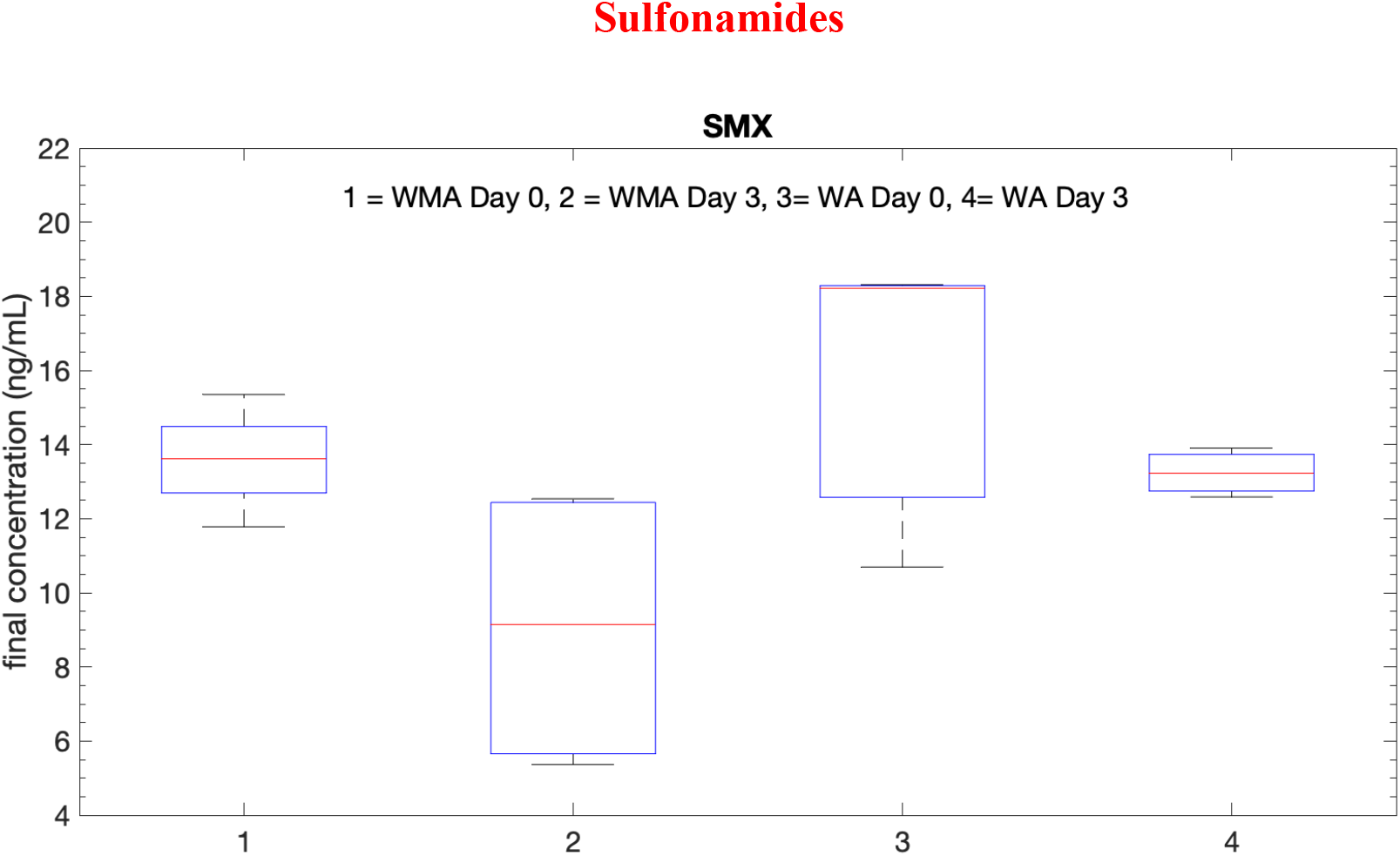
Boxplots for SMX tests.

**Figure 7S.**
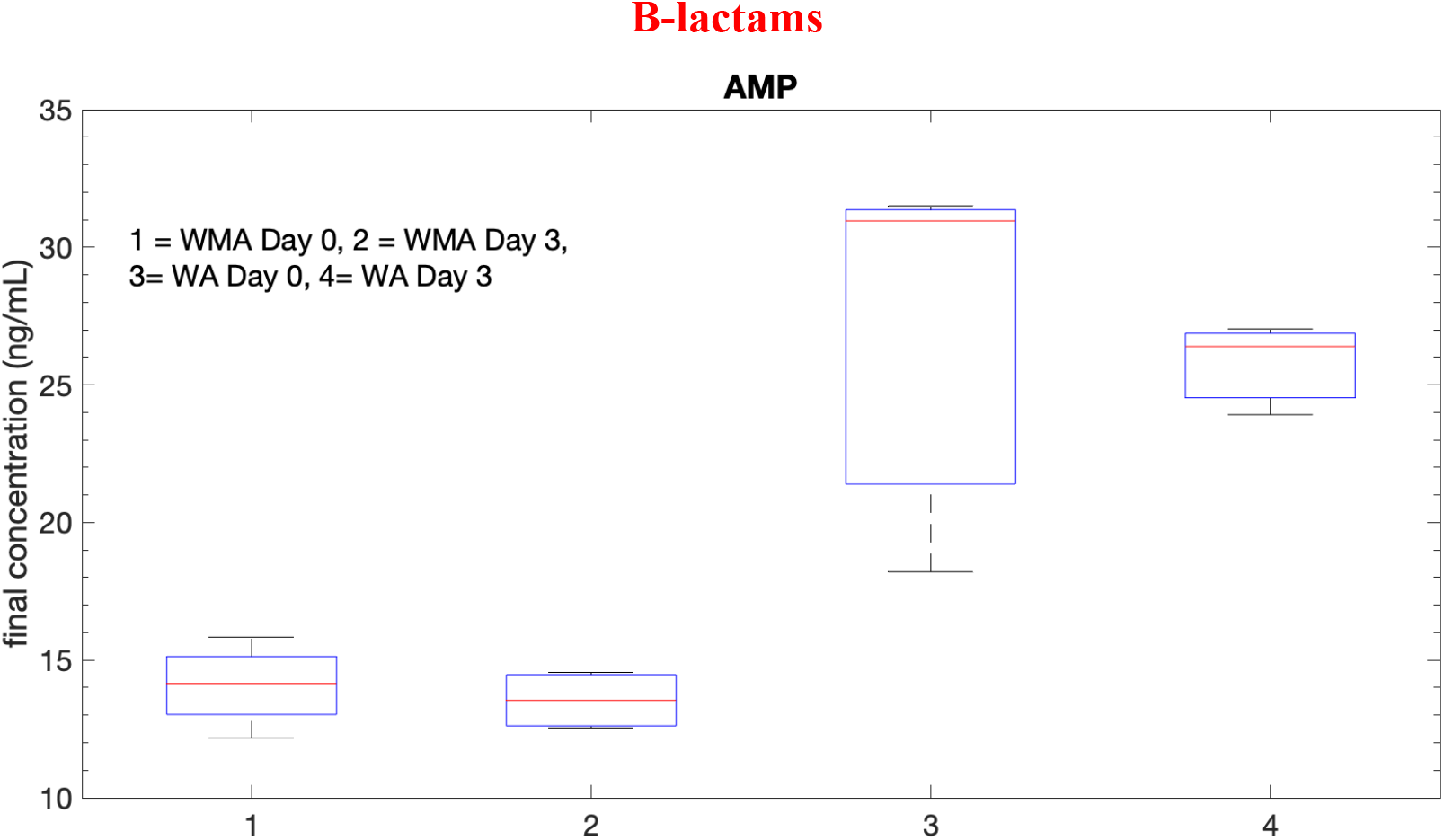
Boxplots for AMP tests.

**Figure 8S.**
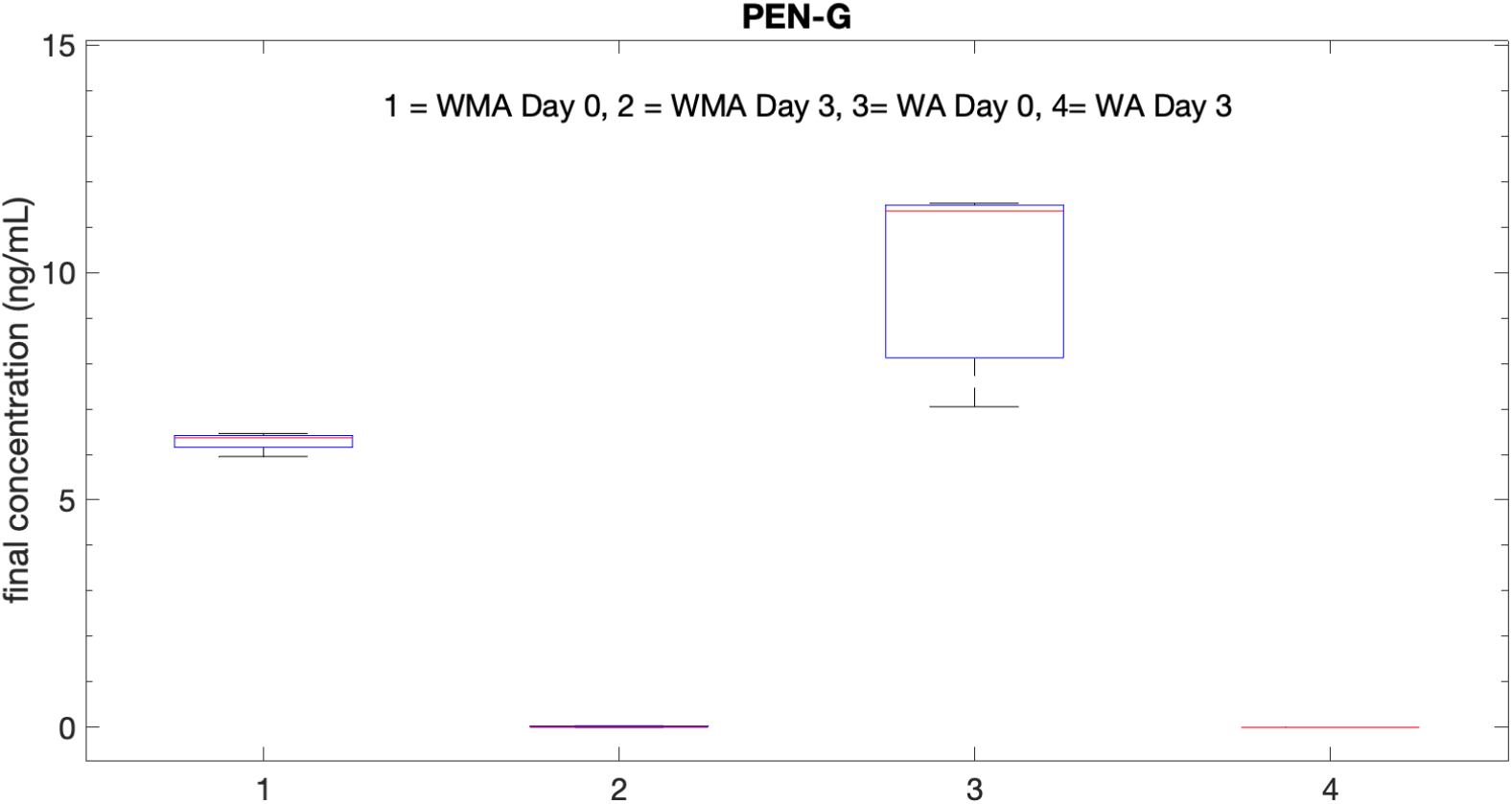
Boxplots for PEN-G tests.

**Figure 9S.**
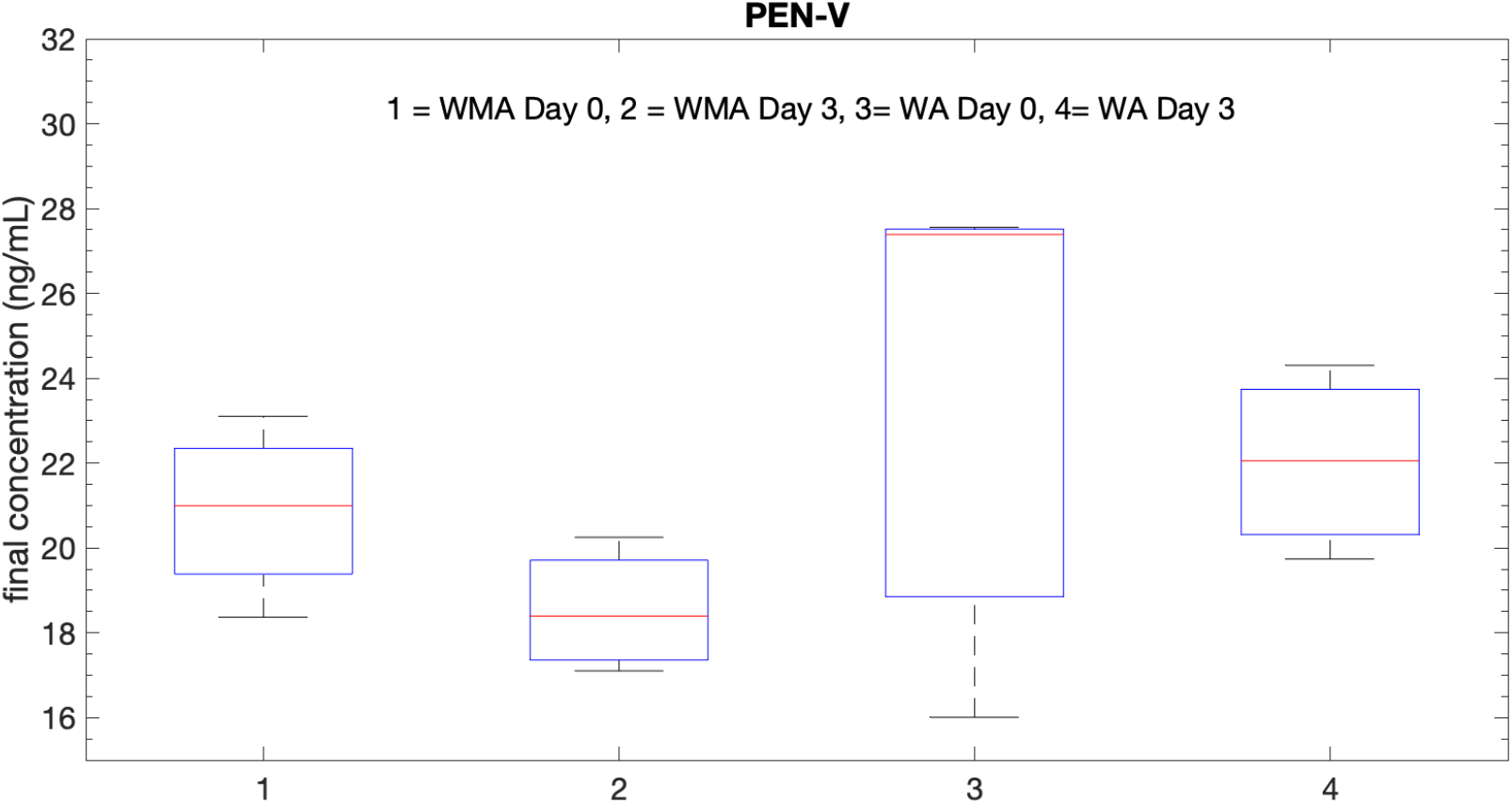
Boxplots for PEN-V tests.

**Figure 10S.**
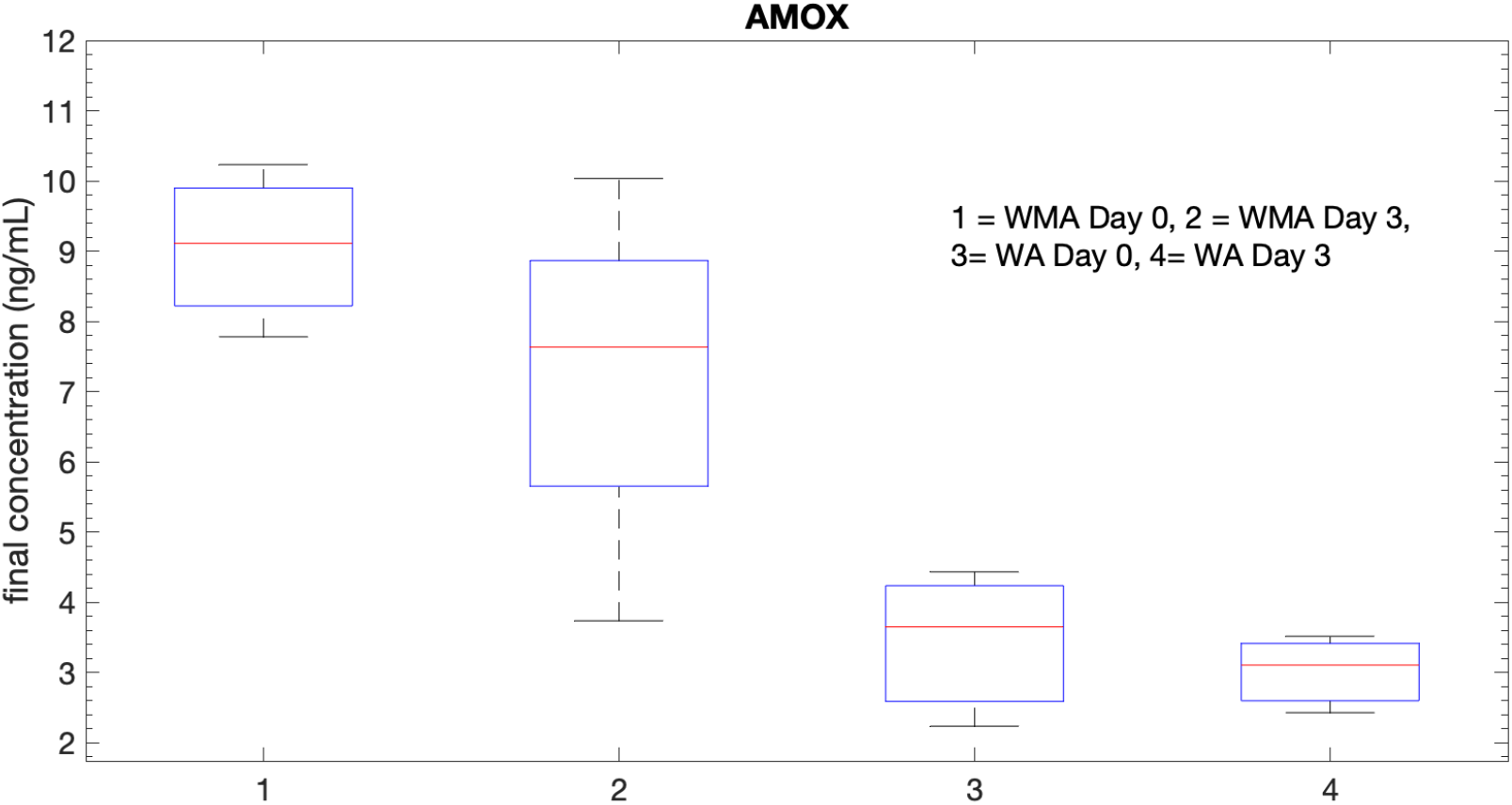
Boxplots for AMOX tests.

**Figure 11S.**
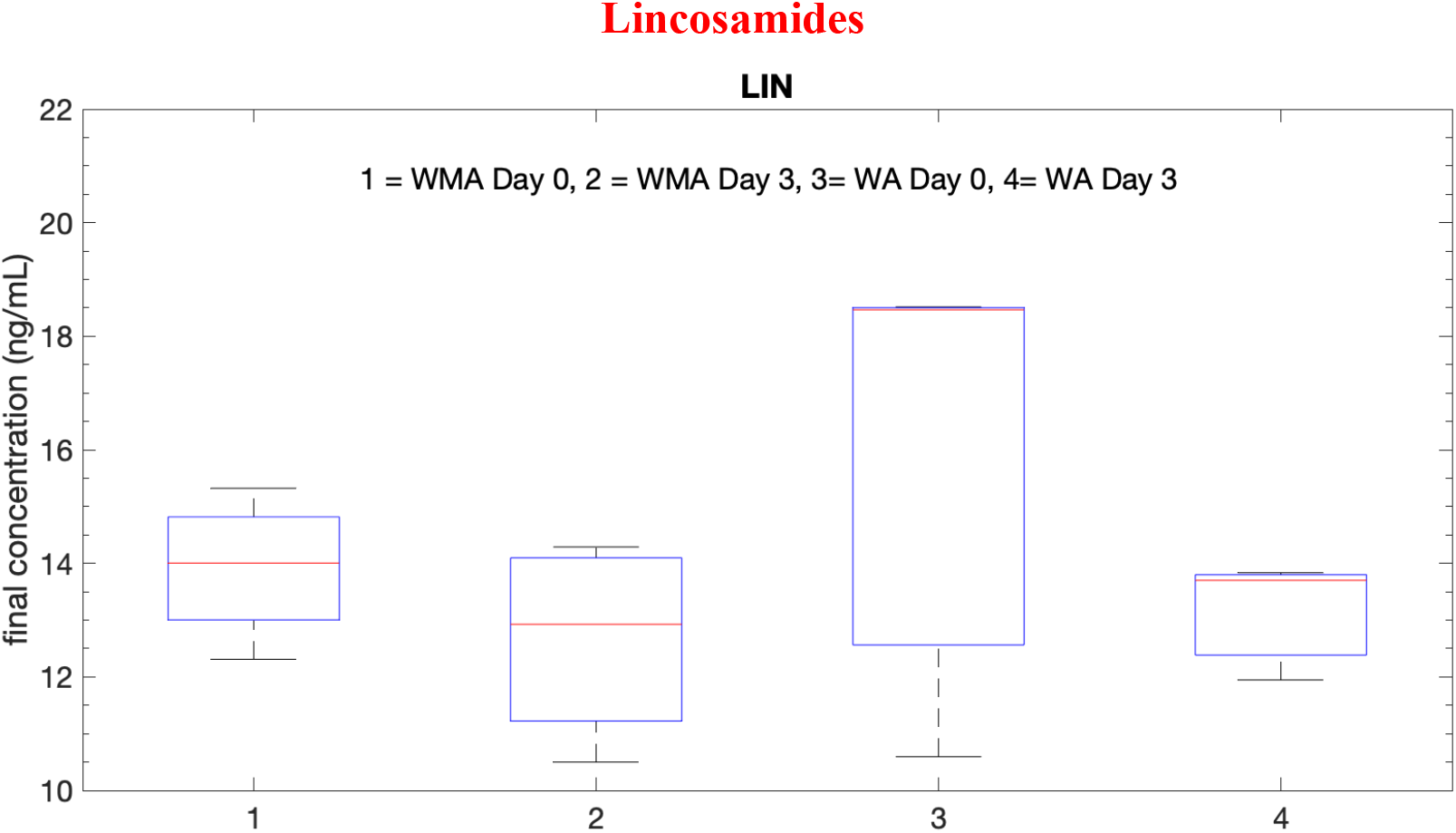
Boxplots for LIN tests.

**Figure 12S.**
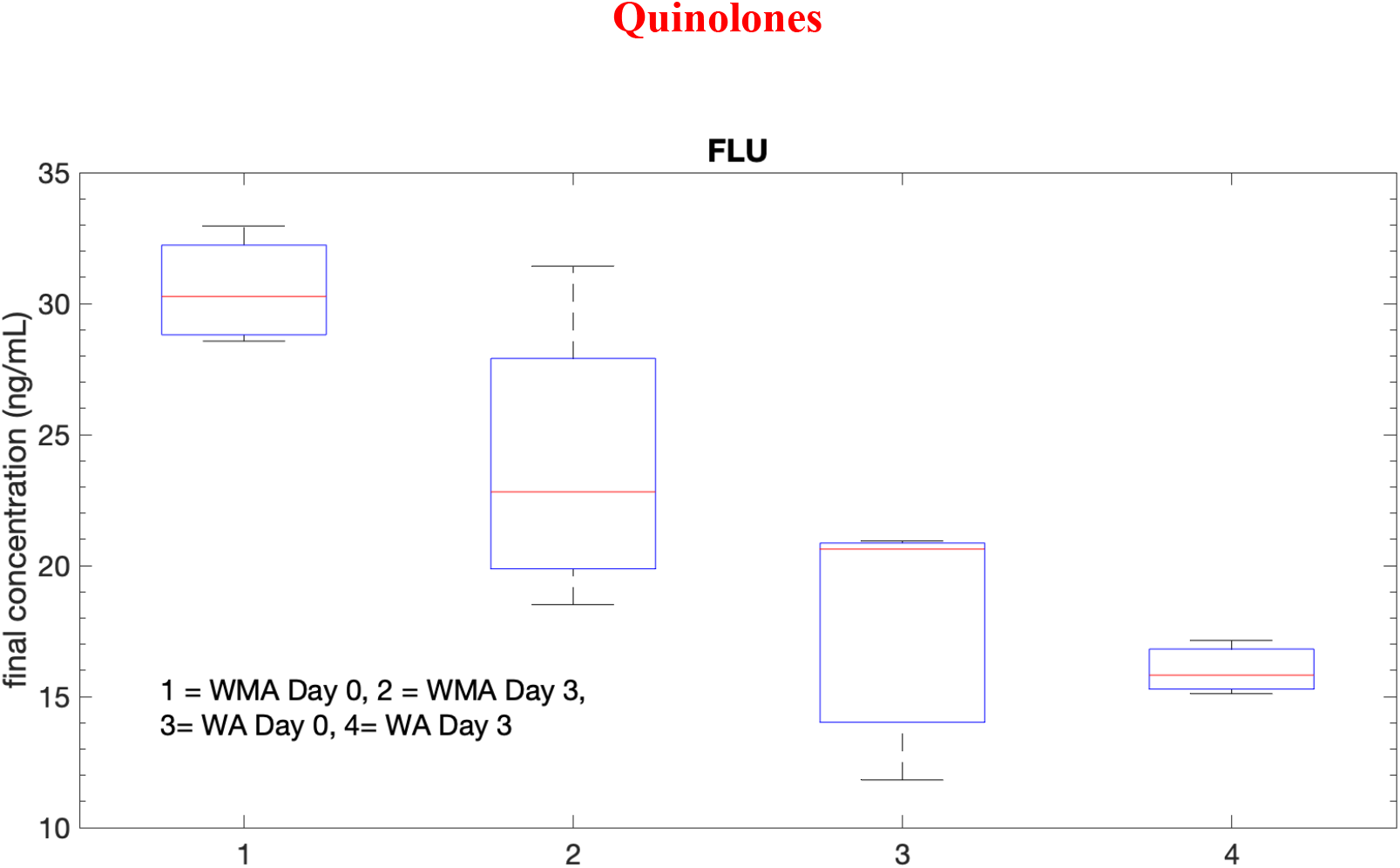
Boxplots for FLU tests.

**Figure 13S.**
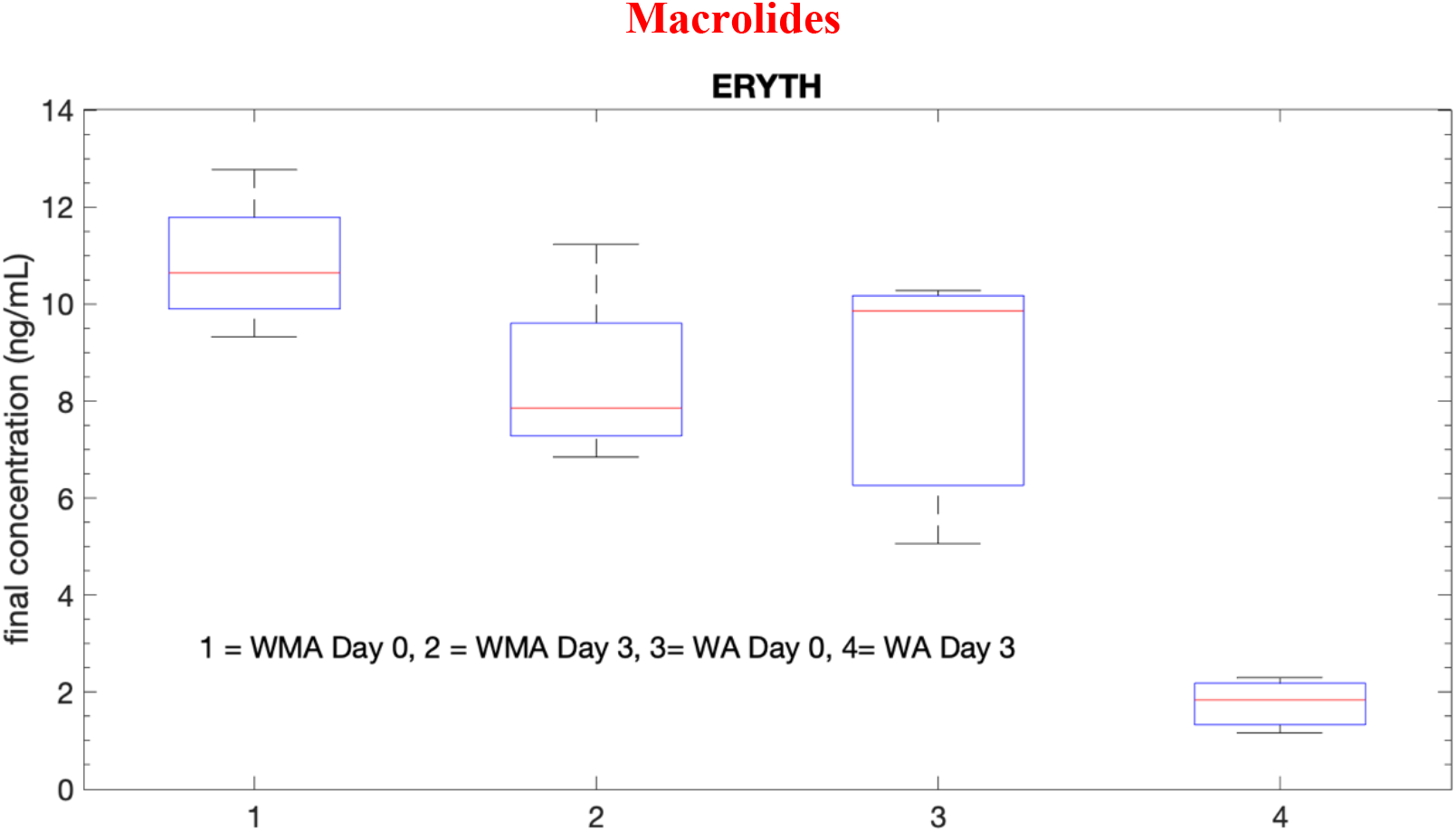
Boxplots for ERYTH tests.

**Figure 14S.**
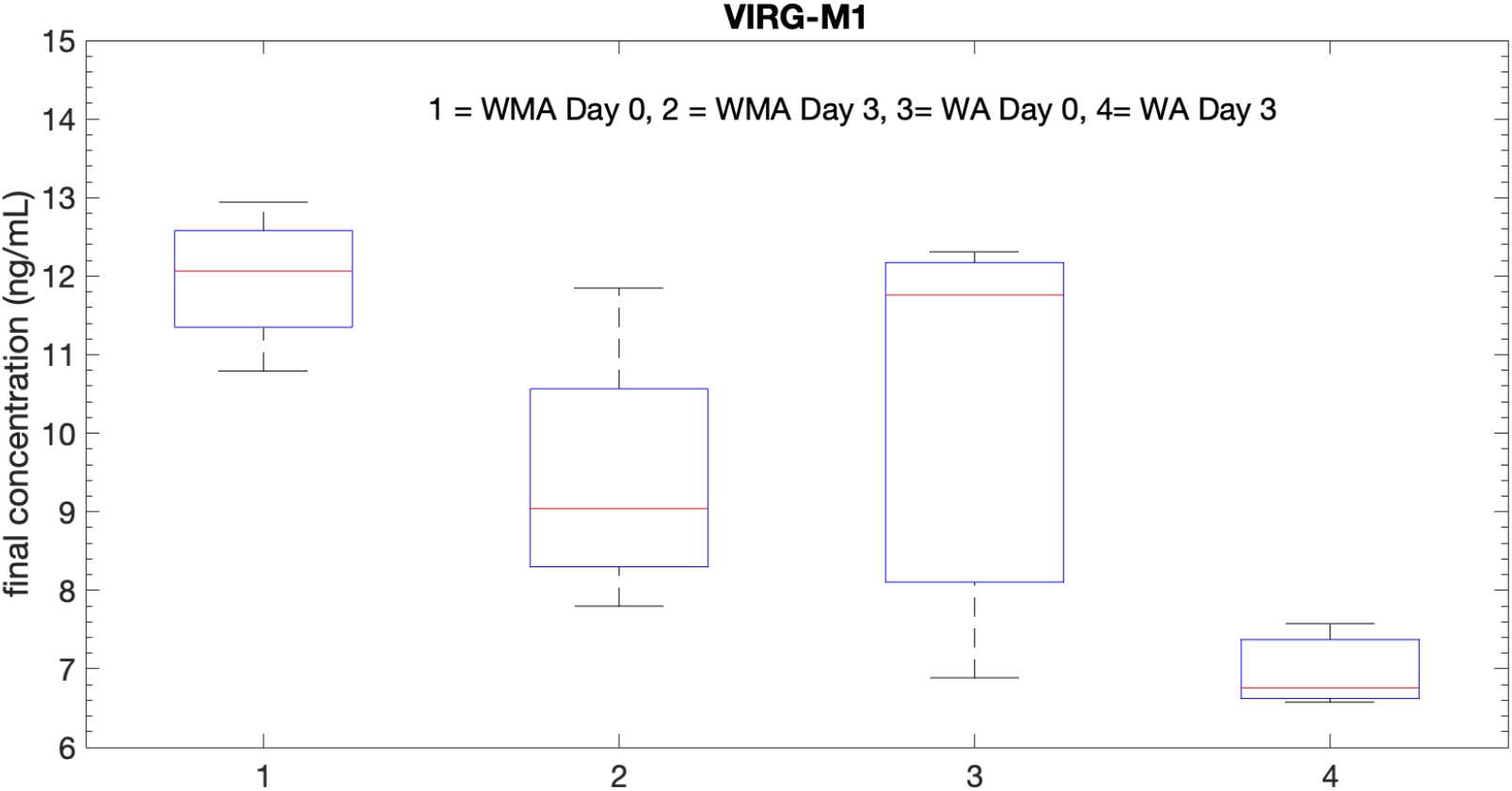
Boxplots for VIRG-M1 tests.

**Figure 15S.**
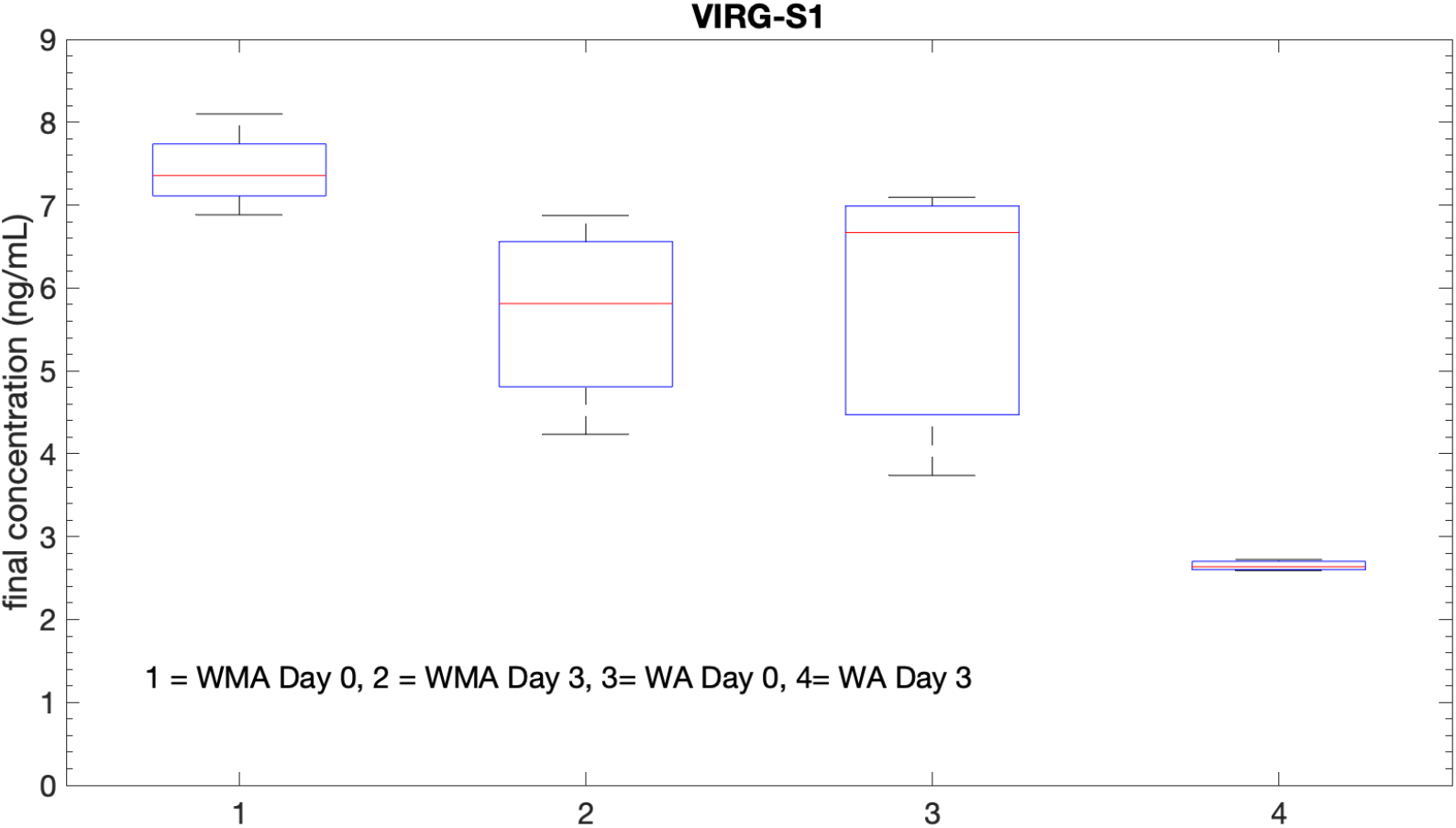
Boxplots for VIRG-S1 tests.

